# ADAPTS: Automated Deconvolution Augmentation of Profiles for Tissue Specific cells

**DOI:** 10.1101/633958

**Authors:** Samuel A Danziger, David L Gibbs, Ilya Shmulevich, Mark McConnell, Matthew WB Trotter, Frank Schmitz, David J Reiss, Alexander V Ratushny

## Abstract

Immune cell infiltration of tumors can be an important component for determining patient outcomes, e.g. by inferring immune cell presence by deconvolving gene expression data drawn from a heterogenous mix of cell types. One particularly powerful family of deconvolution techniques uses signature matrices of genes that uniquely identify each cell type as determined from cell type purified gene expression data. Many methods of this type have been recently published, often including new signature matrices appropriate for a single purpose, such as investigating a specific type of tumor. The package **ADAPTS** helps users make the most of this expanding knowledge base by introducing a framework for cell type deconvolution. **ADAPTS** implements modular tools for customizing signature matrices for new tissue types by adding custom cell types or building new matrices *de novo*, including from single cell RNAseq data. It includes a common interface to several popular deconvolution algorithms that use a signature matrix to estimate the proportion of cell types present in heterogenous samples. **ADAPTS** also implements a novel method for clustering cell types into groups that are hard to distinguish by deconvolution and then re-splitting those clusters using hierarchical deconvolution. We demonstrate that the techniques implemented in **ADAPTS** improve the ability to reconstruct the cell types present in a single cell RNAseq data set in a blind predictive analysis. **ADAPTS** is currently available for use in **R** on CRAN and GitHub.

## Introduction

Determining cell type enrichment from gene expression data is an useful step towards determining tumor immune context [1, 2]. One family of techniques for doing this involves regression with a signature matrix, where each column represents a cell type and each row contains the average gene expression in that cell type [3, 4]. These signature matrices are constructed using gene expression from samples of a purified cell type. Generally, the publicly available versions of these gene expression signature matrices use immune cells purified from peripheral blood. Genes are included in these matrices based on how well they distinguish the constituent cell types. Although examples exist of both general purpose immune signature matrices, e.g. LM22 [5] and Immunostates [6], and more tissue specific ones e.g. M17 [7], these publicly available matrices are most likely not appropriate for all diseases and tissue types. One such example would be multiple myeloma whole bone marrow samples, which pose multiple challenges: both tumor and immune cells are present, immune cells may have different states than in peripheral blood, and non-immune stromal cells such as osteoblasts and adipocytes are expected play an important role in patient outcomes [8].

One straightforward solution to this problem would be to augment a signature matrix by adding cell types without adding any additional genes. For example, one might find purified adipocyte samples in a public gene expression repository and add the average expression for each gene in the matrix to create an adipocyte augmented signature matrix. While this might work, one might reasonably expect adipocytes to best be identified by genes that are different from those that best characterize leukocytes. Furthermore, it will be unclear which deconvolution algorithm would be most appropriate for applying this new signature matrix to samples. Once cell types have been deconvolved, it will also be unclear which cell types are likely to be confused due to a common lineage or other factors and what to do about that confusion. These problems are exacerbated by newly available single cell RNAseq data, which promises to identify the cell types that are present in a particular sample and gene expression for those cell types, but is hampered by clustering techniques that may incorrectly identify groups of cells as distinct cell types.

We have developed the **R** package **ADAPTS** (Automated Deconvolution Augmentation of Profiles for Tissue Specific cells) to help solve these problems. **ADAPTS** is currently available on CRAN (https://cran.r-project.org/web/packages/ADAPTS) and GitHub (https://github.com/sdanzige/ADAPTS). As the package vignettes already provide step-by-step instructions for applying **ADAPTS** to the aforementioned problems, this manuscript is intended to compliment the package by providing a theoretical understandinf of the **ADAPTS** methodology.

## Materials and Methods

**ADAPTS** aids deconvolution techniques that use a signature matrix, here denoted as *S*, where each column represents a cell type and each row contains the average gene expression in that cell type [3, 4]. These signature matrices are constructed using gene expression from samples of purified cell types, *P*, and include genes that are good for identifying cells of type *c* where *c* ∈ *C* and *C* is a population of cell types to look for in a sample.

Deconvolution estimates the relative frequency of cell types in a matrix of new samples *X* where each column is a sample and each row is a gene expression measurement according to Eq 1.

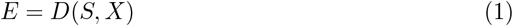

Eq 1 results in a cell type estimate matrix *E*, where each column is a sample corresponding to a column in *X*, and each row is a cell type corresponding to a column in *S*.

One straightforward method to augment a signature matrix, *S*, would be to add new cell types, *NC*, without adding any additional genes. For example, one might start with LM22 as an initial signature matrix, *S*^0^, with |*g*_*S*^0^_ | = 547 genes (rows) and |*C* = 22| cell types (columns) and augment with *c* ∈ *NC* purified cell types. Let *NC*_1_ = adipocytes and *P*^1^ be an adipocyte samples matrix with |*G*| = 20, 000 genes (rows) and |*J*_1_| =9 samples (columns) taken from a public gene expression repository such as ArrayExpress [9] or the Gene Expression Omnibus [10]. A new column could be constructed from *P*^1^ from the average expression, *A*(*P*^1^), for each of the 547 genes (*g*_1_…*g*_547_) in *G*_*S*^0^_. Extended to all *c* ∈ *NC*, this would produce Eq 2.

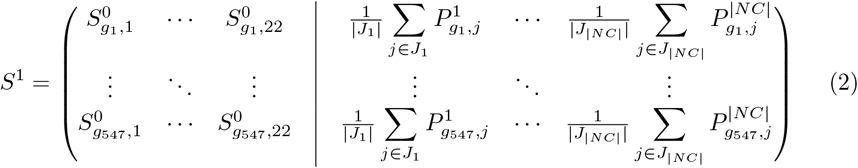

Thus *S*^1^ is a signature matrix augmented with the cell types in NC. While this might work, one might reasonably expect adipocytes to best be identified by genes that are different from those that best characterize the 22 cell types in *S*^0^.

### Signature Matrix Augmentation

**ADAPTS** provides functionality for augmenting an existing cell type signature matrix with additional genes or even constructing a new signature matrix *de novo*. In addition to *S*^0^ and *P*^1^, this requires *S*^*E*0^, an extended version *S*^0^ with all genes. From this data, **ADAPTS** selected *N* additional genes *g*_*n*_1__…*N* to augment the signature matrix as shown in Eq 3.

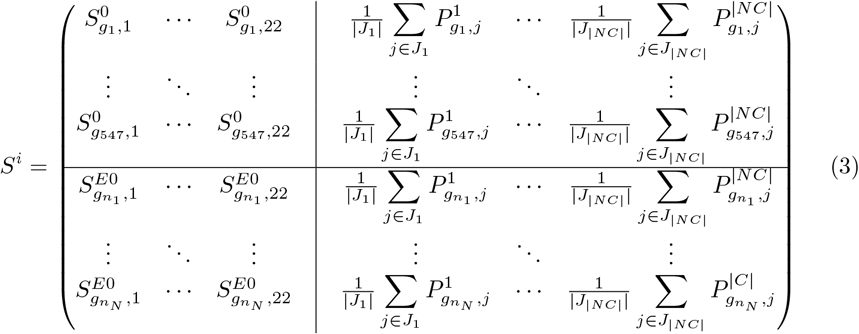

**ADAPTS** helps a user construct new signature matrices with modular **R** functions and default parameters to:

1. Identify and rank significantly different genes for each cell type.
2. Evaluate the stability (condition number, *κ*(*S^x^*)) of many signature matrices *S^x^* ∈ *S*.
3. Smooth and normalize to meet tolerances for a robust signature matrix.

These components are combined into a single function that produces a new deconvolution matrix. First the algorithm ranks each the genes that best differentiate each cell types such that there is a ranked set of genes *g^c^* for each *c* ∈ *C* where *C* includes the cell types in the original signature matrix, *S*^0^ as well as the new cell types *NC*. Genes, *g^c^* (where *g^c^* ⊆ *G* and *G* is the set of all genes), are ranked in descending order according to scores calculated by Eq 4 and exclude any that do not pass a t-test determined false discover rate cutoff (by default, 0.3).

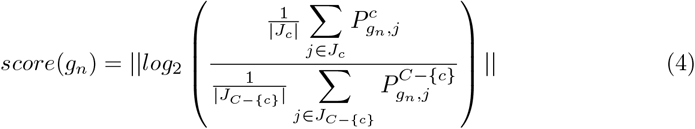

Thus 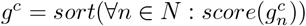 and the function *pop*(*g^c^*) will return and remove the gene with the largest absolute average log expression ratio between the cell type, *c*, and all other cell types, *C* –{*c*}. As shown in Algorithm 1, the matrix augmentation algorithm iteratively adds the top gene that is not already in the signature matrix from each *c* ∈ *C* and calculates the condition number for that matrix. The augmented signature matrix is then chosen that minimizes the condition number, *κ*.

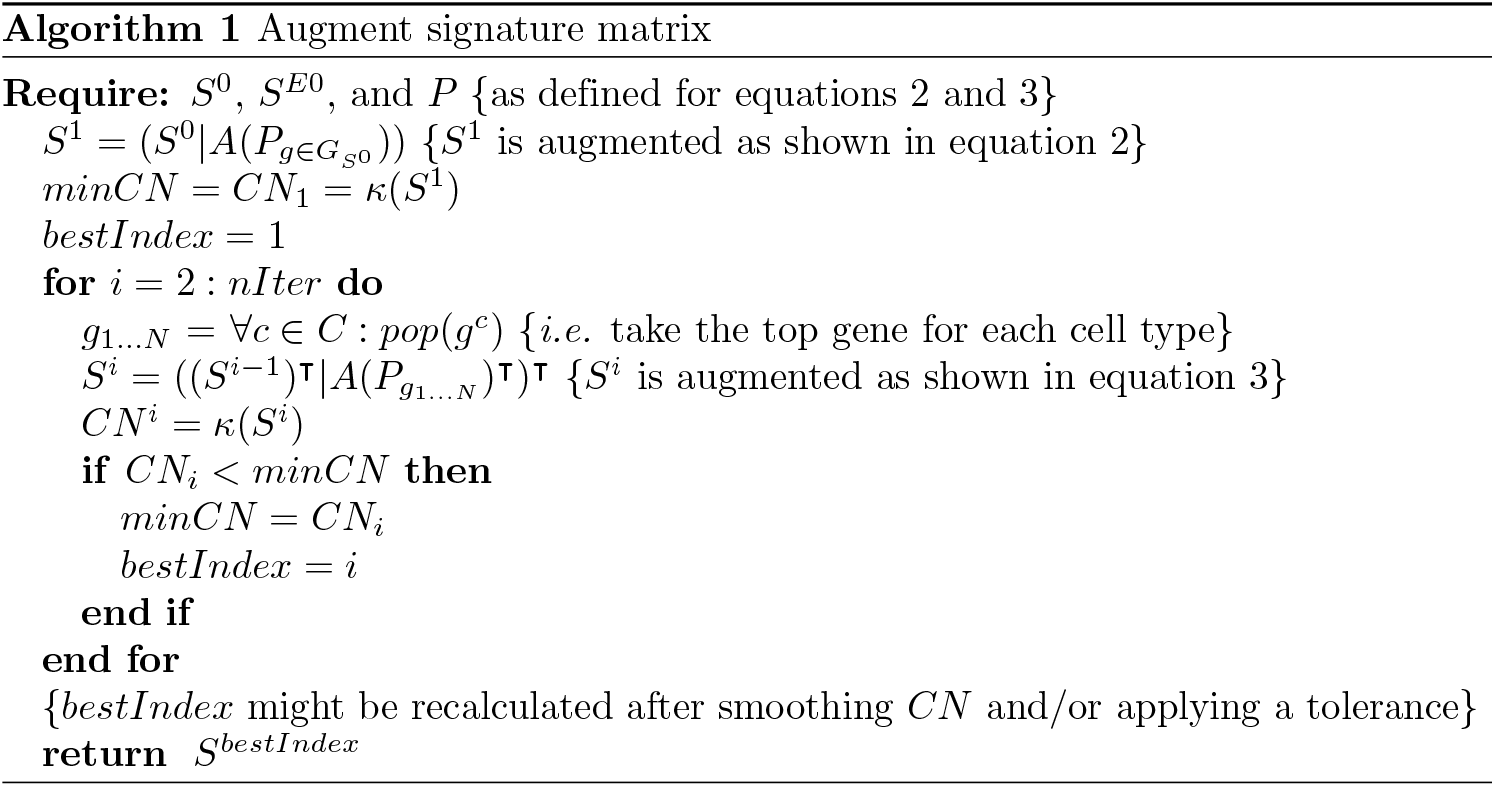

In Algorithm 1: *nIter* = 100 by default, *κ*(*s*) returns the condition number, and *A*(*P*) returns the mean expression for each gene in each cell type. Optionally, the condition numbers (*CN*) may be smoothed to ensure a robust minimum. A tolerance may also be applied to find the minimum number of genes that has a *CN* within some % tolerance of the true minimum.

Fig 1 shows a plot of condition numbers when adding 5 cell types to a 22 cell type signature matrix with smoothing and a 1% tolerance.

**Fig 1.**
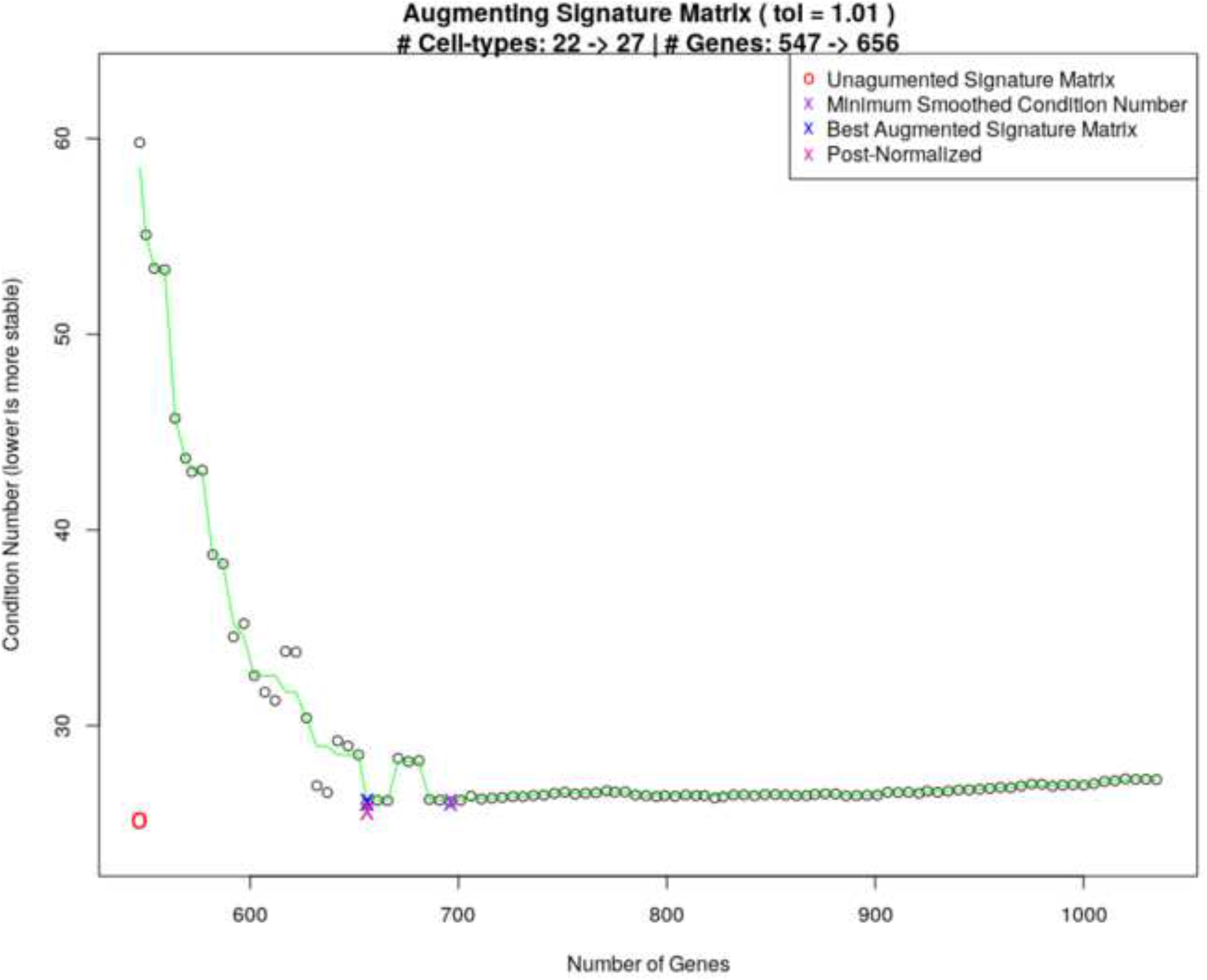
MGSM27 Construction. Curve showing the selection of an optimal condition number for MGSM27.

Similarly, **ADAPTS** can be used to construct a *de novo* matrix from first principals rather than starting with a pre-calculated *S*^0^. One technique is to build *S*^0^ out of the n (e.g. 100) genes that vary the most between cell types and use **ADAPTS** to augment that seed matrix. The n initial genes can then be removed from the resulting signature matrix and that new signature matrix can be re-augmented by **ADAPTS**.

### Deconvolution Framework

The **ADAPTS** package includes functionality to call several different deconvolution methods using a common interface, thereby allowing a user to test new signature matrices with multiple algorithms. These function calls fit the form *D*(*S, X*) presented in Eq 1.

The algorithms include:

1. **DCQ** [11]: An elastic net based deconvolution algorithm that consistently best identifies cell proportions.
2. **SVMDECON** [5]: A support vector machine based deconvolution algorithm.
3. **DeconRNASeq** [12]: A non-negative decomposition based deconvolution algorithm.
4. **Proportions in Admixture** [13]: A linear regression based deconvolution algorithm.

### Spillover to Convergence

In cell-type deconvolution, spillover refers to the tendency of some cell types to be misclassified as other cell types [14]. For example, when using LM22, deconvolving purified activated mast cell samples results predicted cell compositions that are almost equally split between activated and resting mast cells (Figure 2). One approach to exploring this problem might be to cluster the signature matrix, and assume that highly correlated signatures would tend to spill over to each other. However, **ADAPTS** instead directly calculate what cell types spill over to what other cell types by deconvolving the purified samples, *P*, used to construct and augment the signature matrices, *S*. While the cell types that are likely to spill-over detected by both methods are similar, directly calculating the spillover reveals some surprising patterns. For example, based on signature matrix clustering of LM22, ‘Dendritic.cells.activated’ and ‘Dendritic.cells.resting’ tend to cluster together, however the spillover patterns (Figure 2) reveal that ‘Dendritic.cells.activated’ are most similar to ‘Macrophages.M1’ while ‘Dendritic.cells.resting’ are similar to ‘Macrophages.M1’ and ‘Macrophages.M2’.

**Fig 2.**
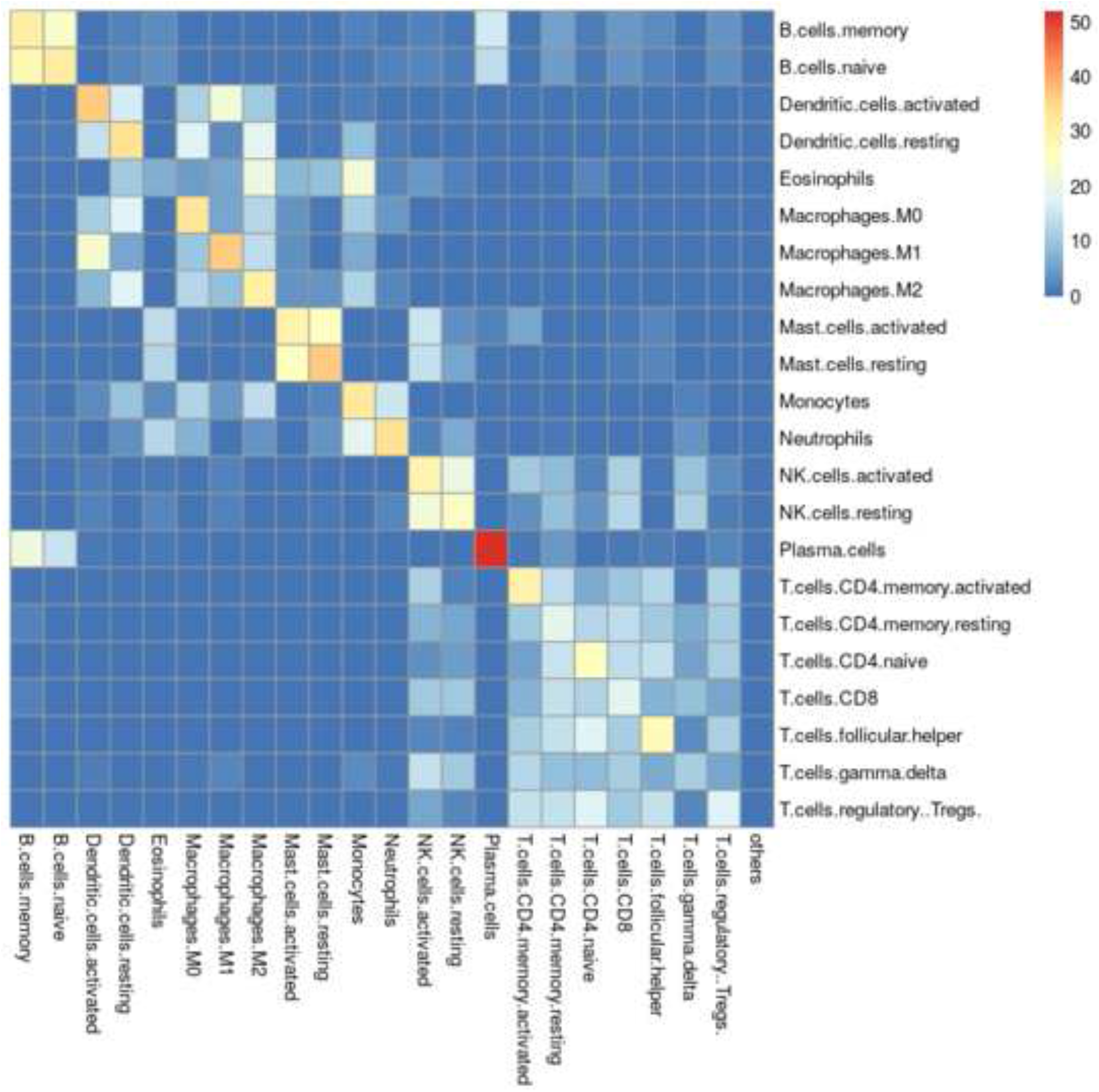
LM22 Spillover Matrix. Spillover matrix showing mean misclassification of purified samples for LM22. Rows show purified cell types and columns show what those samples are deconvolved as.

As shown in Algorithm 2, recursively (or iteratively) applying the spillover calculation reveals clear clusters of cells. Eq 5 revisits Eq 1, obtaining an initial spillover matrix, *E*^0^, by applying Eq 1 to a signature matrix, *S*^0^, and the source data used to construct it, *P*^0^.

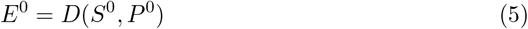

Applying *A*(*P*) to average the cell type estimates *E* across purified samples makes the spillover matrix resemble a signature matrix, leading to Eq 6.

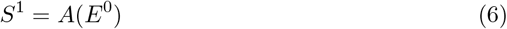

This new spillover based deconvolution matrix *S*^1^ can be used to re-deconvolve the initial spillover matrix, *E*^0^, effectively ‘sharpening’ the deconvolution matrix image as shown in Eq 7.

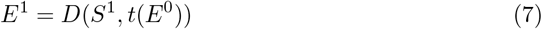

Once these values are calculated, the following pseudocode (Algorithm 2) shows how **ADAPTS** iteratively applies spillover re-deconvolution to cluster cell types likely to be confused by deconvolution.

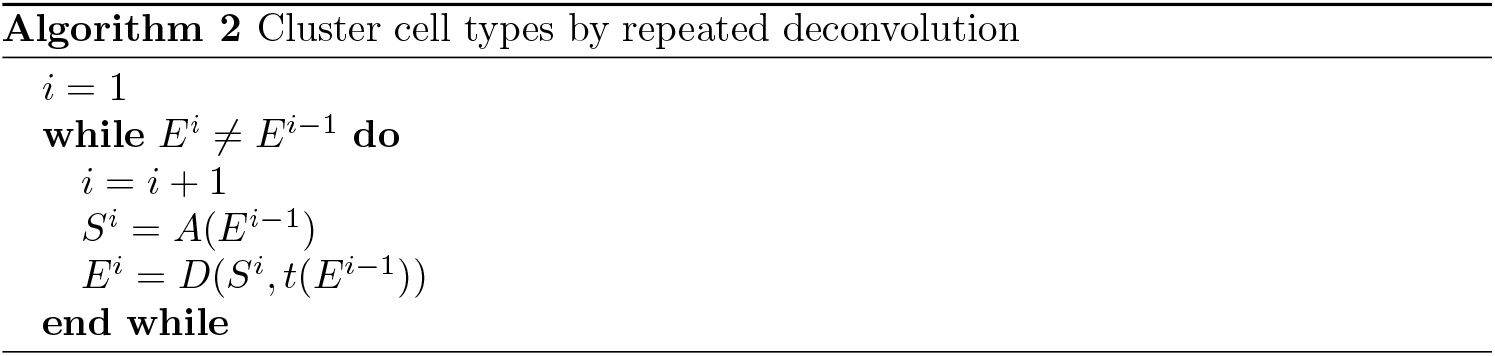

As shown in Algorithm 2, the signature matrix may never converge onto a single question, but instead may alternate between several solutions such that *E^i^* = *E*^*i*–1^ is impossible. Therefore **ADAPTS** includes a parameter forcing the algorithm to break and return an answer after *i* iterations. However, the algorithm usually converges in less that 30 iterations, resulting in a clustered spillover matrix (*e.g*. Fig 3).

**Fig 3.**
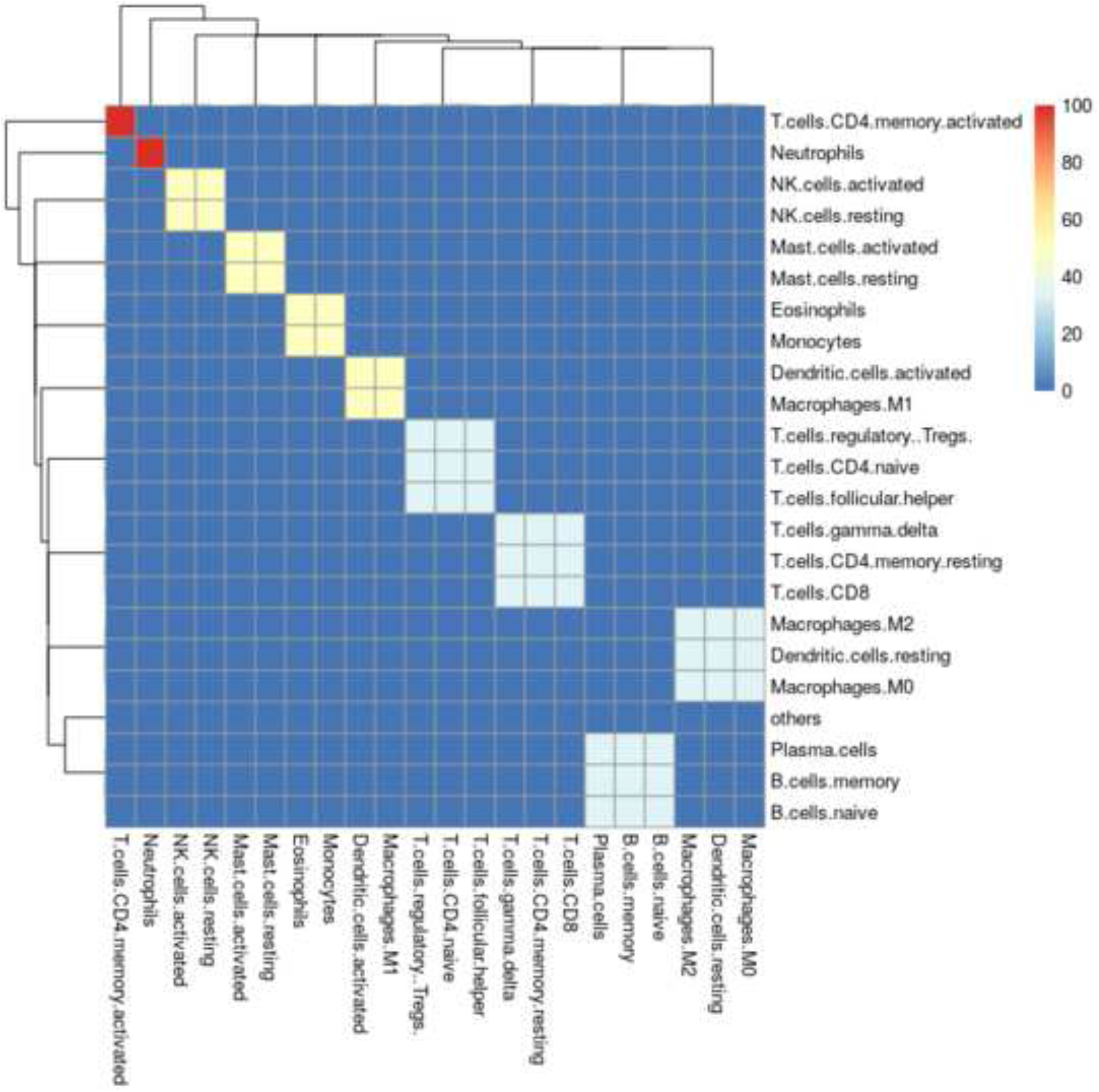
LM22 converged spillover matrix. Iterative deconvolution shows how easily confused cell types conspicuously form clusters.

The resulting cell-type clusters (*CC*) are extracted from *E^i^* by grouping the cell-types for any rows that are identical. For example in Fig 3, ‘NK.cells.activated’ and ‘NK.cells.resting’ would be grouped in one cluster (e.g. *CC*^3^), while ‘Neutrophils’ would exist in a cluster by themselves (*CC*^2^), and |*CC*| = 10.

### Hierarchical Deconvolution

The clusters calculated by Algorithm 2 allow the hierarchical deconvolution implemented in **ADAPTS. ADAPTS** includes a function to automatically train deconvolution matrices that include only genes that differentiate cell types that cluster together in Algorithm 2. The first round of deconvolution determines the total fraction of cells in the cluster. The next round of deconvolution determines the relative proportion of all of the cell types in that cluster as shown in Algorithm 3.

While this algorithm has not been implemented recursively in **ADAPTS**, if it was it would resemble a discrete version of the continuous model implemented in MuSiC [15].

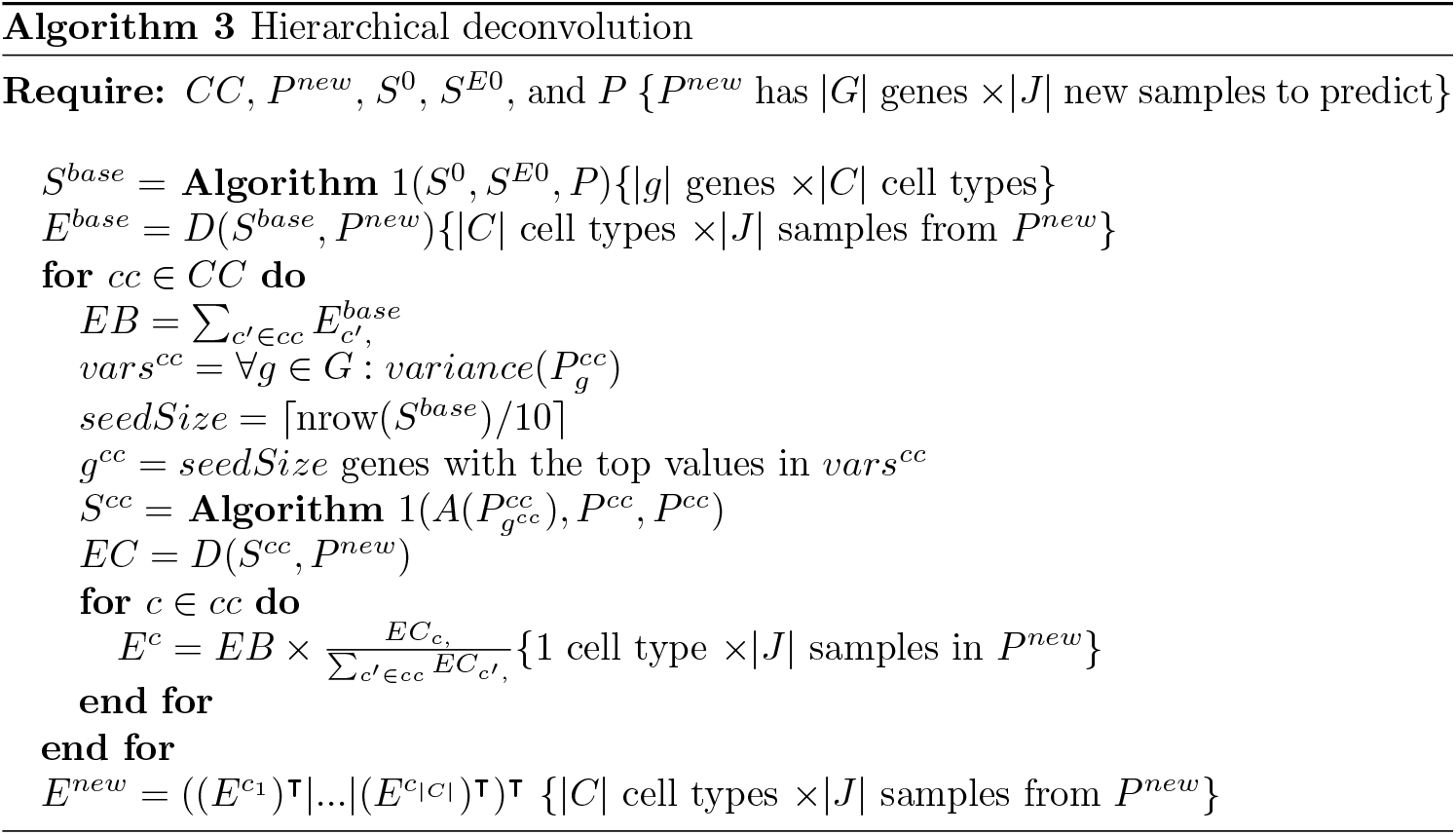

## Results

The following results section shows how the theory set out in Materials and Methods is applied to detect tumor cells in multiple myeloma samples and to utilize single cell RNAseq data to build a new signature matrix. It contains highlights from two vignettes distributed with the CRAN package (S1 Vig and S2 Vig).

### Example: Detecting Tumor Cells

To demonstrate utility of the **ADAPTS** package, we show how it can be used to augment the LM22 from [5] to identify myelomatous plasma cells from gene expression profiles of 423 purified tumor (CD138^+^) samples and 440 whole bone marrow (WBM) samples taken from multiple myeloma patients. The fraction of myeloma cells, which are tumorous plasma cells, were identified in both sample types via quantification of the cell surface marker CD138. Root mean squared error (RMSE) and Pearson’s correlation coefficient (*ρ*) were used to evaluate accuracy of tumor cell fraction estimates. RMSE proved particularly relevant when deconvolving purified CD138^+^ sample profiles, because 356 of 423 samples are more than 90% pure tumor resulting in clumping of samples with purity near 100%.

The following matrices were used or generated during the evaluation:

- **LM22:** As reported in [5]. The sum of the ‘memory B cells’ and ‘plasma cells’ deconvolved estimates represent tumor percentage.
- **LM22 + 5:** Builds on LM22 by adding purified sample profiles for myeloma specific cell types as shown in Eq 2: plasma memory cells [16], osteoblasts [9], osteoclasts, adipocytes, and myeloma plasma cells [17]. The sum estimates for ‘memory B cells’, ‘myeloma plasma cells’, ‘plasma cells’, and ‘plasma memory cells’ represent tumor percentage.
- **MGSM27:** Builds on LM22 by adding 5 myeloma specific cell types using **ADAPTS** to determine inclusion of additional genes as shown in Eq 3. Fig 1 shows **ADAPTS** evaluating matrix stability after adding different numbers of genes, smoothing the condition numbers, and selecting an optimal number of features.
- *de novo* **MGSM27**: Builds a *de novo* MGSM27 by seeding with the 100 most variable genes from publicly available data similar to those mentioned in [5] and the 5 aforementioned myeloma specific cell types.

Table 1 displays average *RMSE* and *ρ* for tumor fraction estimates obtained via application of DCQ deconvolution using the four aforementioned matrices across both myeloma profiling datasets.

**Table 1.**
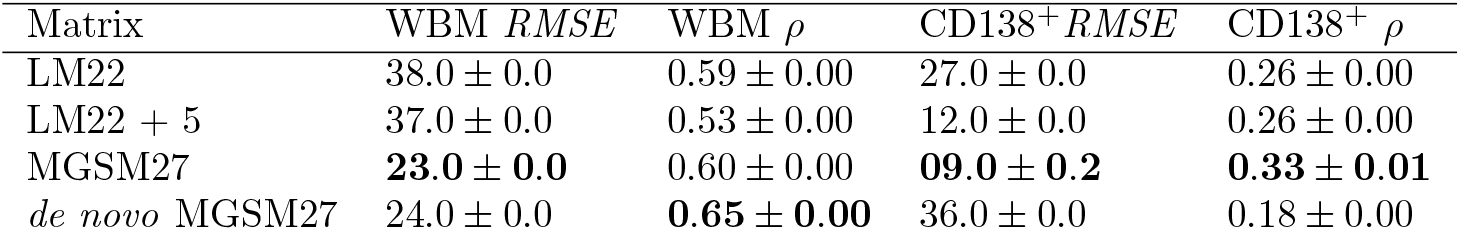
Deconvolution reconstruction of tumor percentage in whole bone marrow (*WBM*) and samples sorted to consist of nearly pure CD138^+^ cells. Classifier accuracy is measured by root mean square error (*RMSE*) and Pearson’s correlation coefficient (*ρ*). The best scores in each column are bolded.

| Matrix | WBM $RMSE$ | WBM $\rho$ | CD138<sup>+</sup> $RMSE$ | CD138<sup>+</sup> $\rho$ |
| --- | --- | --- | --- | --- |
| LM22 | 38.0 ± 0.0 | 0.59 ± 0.00 | 27.0 ± 0.0 | 0.26 ± 0.00 |
| LM22 + 5 | 37.0 ± 0.0 | 0.53 ± 0.00 | 12.0 ± 0.0 | 0.26 ± 0.00 |
| MGSM27 | **23.0 ± 0.0** | 0.60 ± 0.00 | **09.0 ± 0.2** | **0.33 ± 0.01** |
| *de novo* MGSM27 | 24.0 ± 0.0 | **0.65 ± 0.00** | 36.0 ± 0.0 | 0.18 ± 0.00 |

While the exact genes chosen during each run varies slightly, Table 1 shows that consistently the best accuracy is achieved by augmenting LM22 using **ADAPTS**.The reduced performance of the *de novo* MGSM27 on the *C*D138^+^ samples is likely due to genes that were present in LM22, but were missing in some of the source data and thus excluded from *de novo* construction. More details are available in the vignette distributed with the **R** package.

### Spillover Matrix

Successfully recapturing the known percentage of tumor cells in a sample is a useful intermediate validation step, however, the true value of a deconvolution algorithm lies in it’s ability to determine cell types in a sample that affect patient outcomes. Statistical and machine learning techniques may be applied to identify relevant cell estimates. From there, a correct understanding of the limitations of deconvolution is helpful to reveal the underlying biology. One particularly relevant limitation of deconvolution is how the algorithm may confuse different cell types. **ADAPTS**’s approach to resolving these problems are outlined in and results in plots such as those shown in Figs 2 and 3.

This sort of analysis leads to Algorithm 2 and cell type clusters such as those shown in Fig 3. One way to interpret these results is that co-clustered cell types are those cannot be reliably distinguished by deconvolution using a particular deconvolution algorithm and signature matrix. In this example, ‘B.cells.naive’, ‘B.cells.memory’, and ‘Plasma.cells’ are all clearly clustered together. These clusters may be particularly valuable for single cell RNAseq analysis where clustering software such as Seurat [18] aid in annotating cell types, but can introduce artificial distinctions due to limitations inherent in clustering.

## Example: Deconvolving Single Cell Pancreas Samples

In this section we demonstrate how **ADAPTS** can be applied to build a deconvolution matrix from single cell RNAseq data. This example has the additional benefit of illustrating the utility of the algorithms outlined in Spillover to Convergence and Hierarchical Deconvolution to find cell type clusters and distinguish between cell types in those clusters. In this example we use the pancreas single cell RNAseq dataset available in in Array Express [9] as E-MTAB-5061 [19]. All cells of single type were combined and averaged to build pseudo-pure samples of each annotated cell types. A pseudo-bulk RNAseq sample was constructed by adding together all cell types, with the pseudo-bulk cell type percentages assigned based on the proportion of annotated single cells in the mix. The normal pancreas samples were used as the training set and the diabetic pancreas samples as the test set.

To demonstrate the utility of augmenting a signature matrix with **ADAPTS**, we build a signature matrix from the top 100 most variant genes (i.e. Top100) and then augmented this signature (i.e Augmented) as shown in Fig 4. The first test is to predict the normal pseudo-bulk data - essentially predicting the training set (Table 2). The second test is a blind estimation of the diabetic pancreas sample (table 3). As shown in Table 2 the Top100 genes set the baseline correlation coefficient (i.e. *ρ*) at 0.05 and the root mean square error (*RMSE*) at 13.82. Augmenting the signature matrix with **ADAPTS** Algorithm 1 improved the rho to 0.26 and RMSE to 10.72.

**Fig 4.**
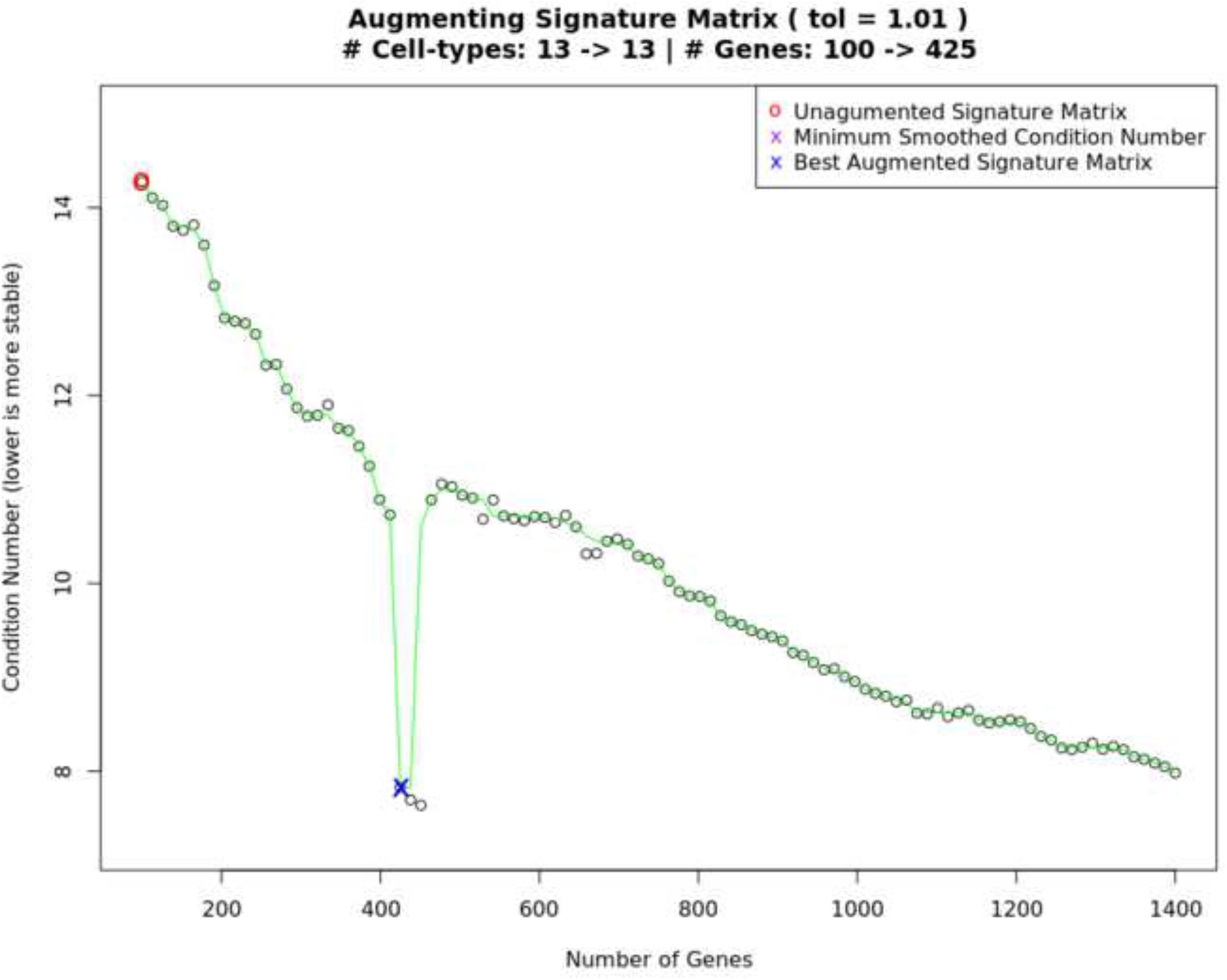
scRNAseq Signature Matrix Construction. Curve showing the selection of an optimal condition number for the single cell RNAseq augmented signature matrix data.

**Table 2.** Deconvolution cell type estimates of the normal pancreas training set.

| Cell Type | Top 100 | Top 100 hierarchical | Augmented | Augmented hierarchical | Reference |
| --- | --- | --- | --- | --- | --- |
| acinar.cell | 11.38 | 7.88 | 11.56 | 10.49 | 10.36 |
| ductal.cell | 7.32 | 7.85 | 7.85 | 12.56 | 43.11 |
| alpha.cell | 7.46 | 7.92 | 8.34 | 9.51 | 12.11 |
| gamma.cell | 11.66 | 7.77 | 9.68 | 8.51 | 1.83 |
| beta.cell | 7.11 | 11.75 | 7.66 | 10.77 | 3.51 |
| co.expression.cell | 4.36 | 16.91 | 11.49 | 8.41 | 14.26 |
| delta.cell | 0.00 | 12.27 | 2.56 | 8.04 | 0.88 |
| unclassified.endocrine.cell | 7.02 | 20.63 | 6.11 | 12.29 | 0.40 |
| endothelial.cell | 8.37 | 0.00 | 7.63 | 0.00 | 8.53 |
| PSC.cell | 0.00 | 0.00 | 2.13 | 8.52 | 0.32 |
| epsilon.cell | 0.00 | 7.02 | 2.68 | 6.11 | 0.16 |
| mast.cell | 0.00 | 0.00 | 5.96 | 0.80 | 2.55 |
| MHC.class.II.cell | 35.33 | 0.00 | 16.37 | 4.01 | 1.99 |
| others | 0.00 | 0.00 | 0.00 | 0.00 | 0.00 |
| *RMSE* | 13.82 | 12.09 | 10.72 | 10.16 | 0.00 |
| $\rho$ | 0.05 | 0.12 | 0.26 | 0.39 | 1.00 |

**Table 3.** Blind deconvolution cell type estimates of the diabetic pancreas test set.

| Cell Type | Top 100 | Top 100<br>hierarchical | Augmented | Augmented<br>hierarchical | Reference |
| --- | --- | --- | --- | --- | --- |
| acinar.cell | 7.40 | 4.32 | 10.82 | 8.89 | 5.78 |
| alpha.cell | 7.93 | 9.01 | 7.94 | 14.41 | 36.24 |
| beta.cell | 8.04 | 8.44 | 8.58 | 9.35 | 12.39 |
| co.expression.cell | 12.33 | 7.95 | 10.03 | 8.55 | 1.68 |
| delta.cell | 9.38 | 11.50 | 8.13 | 10.93 | 7.35 |
| ductal.cell | 5.93 | 17.06 | 12.49 | 8.90 | 21.74 |
| endothelial.cell | 0.00 | 13.80 | 2.58 | 7.91 | 0.53 |
| epsilon.cell | 6.56 | 21.36 | 5.83 | 12.12 | 0.21 |
| gamma.cell | 8.46 | 0.00 | 7.61 | 0.00 | 9.45 |
| mast.cell | 0.00 | 0.00 | 1.72 | 8.51 | 0.32 |
| MHC.class.II.cell | 0.00 | 6.56 | 2.88 | 5.83 | 0.32 |
| PSC.cell | 0.00 | 0.00 | 5.93 | 0.55 | 2.31 |
| unclassified.endocrine.cell | 33.97 | 0.00 | 15.47 | 4.05 | 1.68 |
| others | 0.00 | 0.00 | 0.00 | 0.00 | 0.00 |
| *RMSE* | 12.76 | 10.68 | 9.44 | 8.91 | 0.00 |
| $\rho$ | 0.06 | 0.24 | 0.35 | 0.46 | 1.00 |

### Clustering Cell Types Improves Deconvolution Accuracy

The spillover clustering algorithm outlined in section was applied to the Top100 and Augmented signature matrices. Fig 5 shows the cell type clusters for the Top100 signature matrix, and Fig 6 for the Augmented signature matrix. One way to interpret the results is to assume that the clustered cell-types are indistinguishable from each other, then the correct comparison method is to treat both as the same cell type.

**Fig 5.**
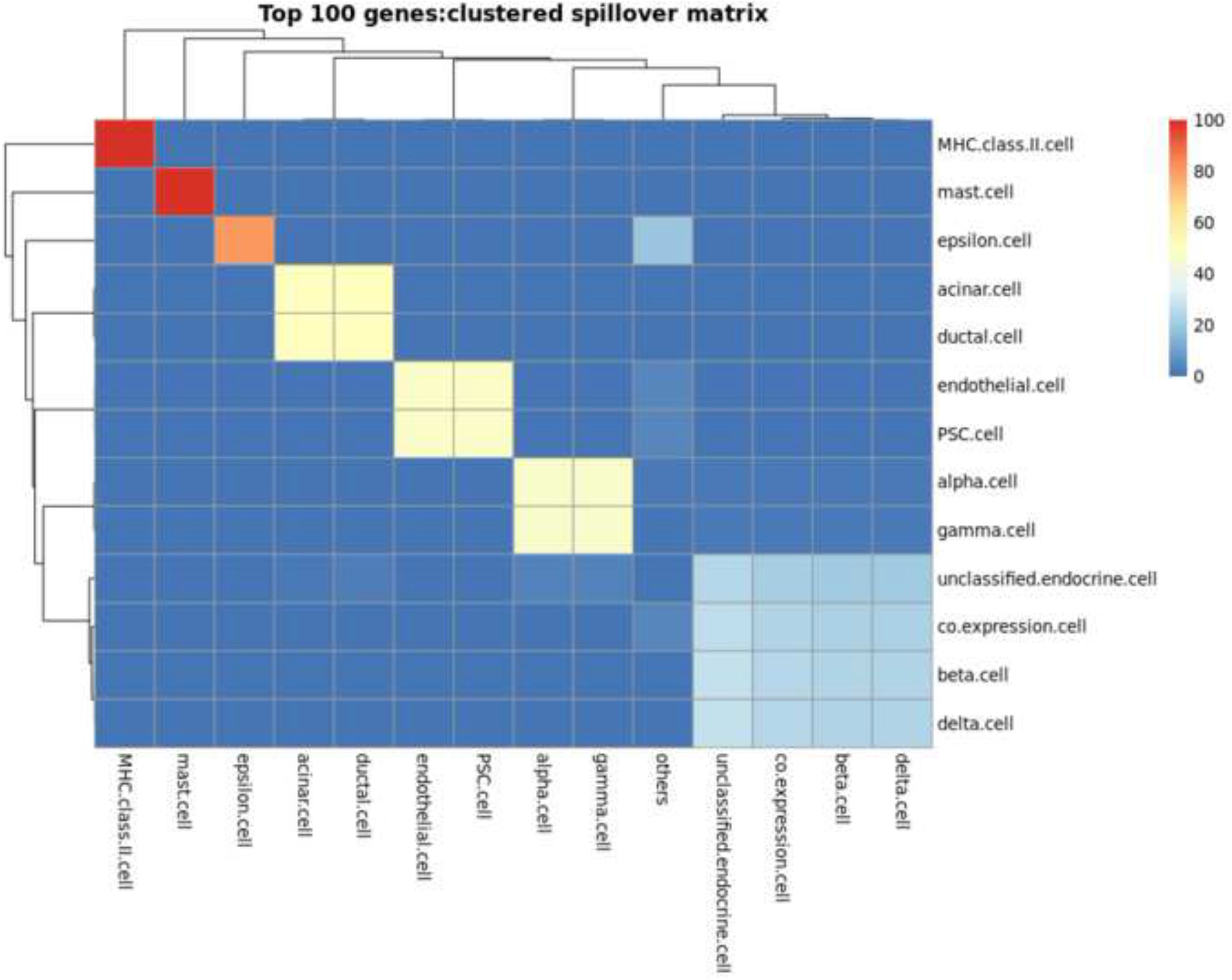
Clustering of Top 100 gene signature matrix. The cell type clusters identified using the signature matrix constructed from the 100 genes with the highest variance across cell types in the single cell data drawn from a normal pancreas sample.

**Fig 6.**
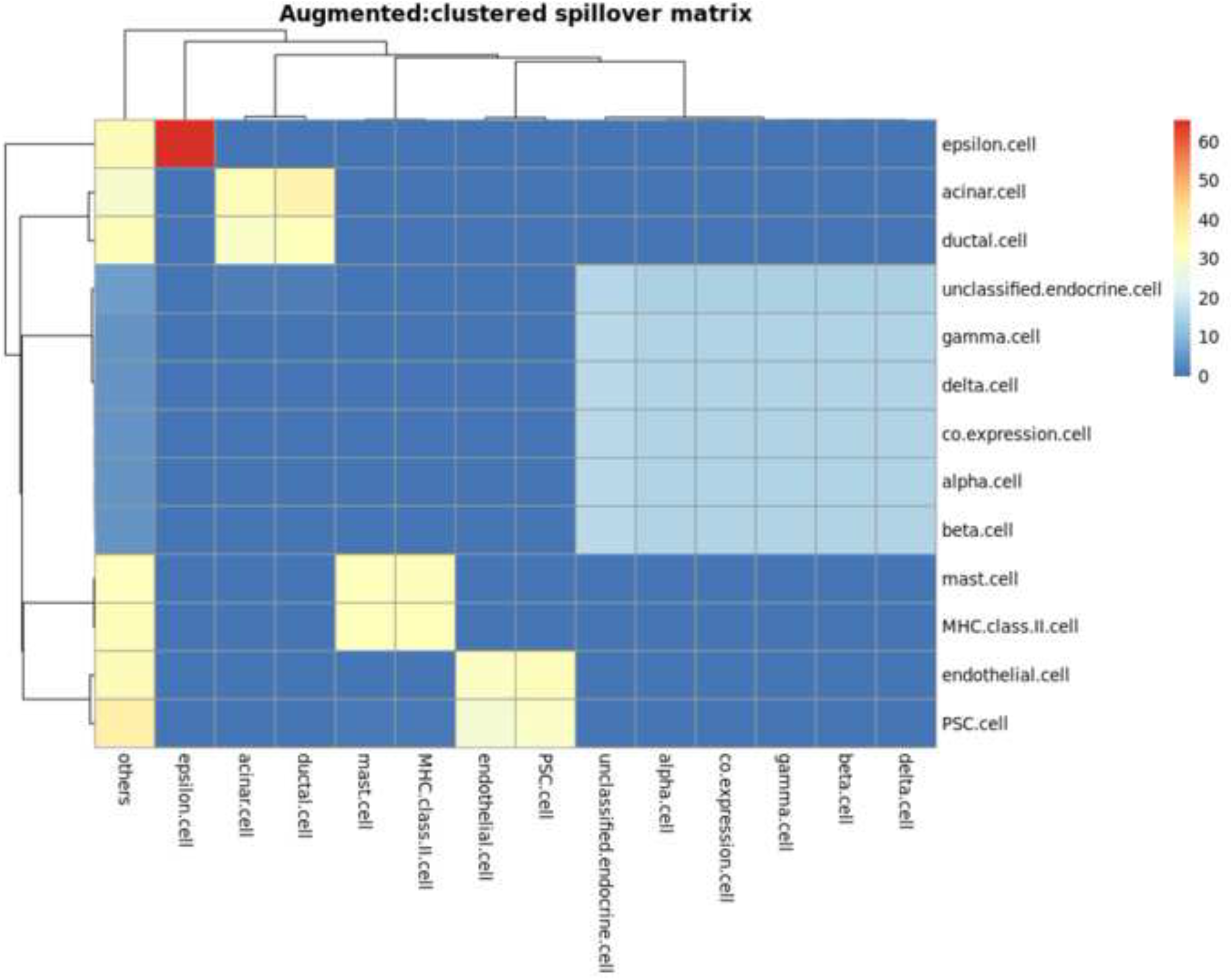
Clustering of Augmented gene signature matrix. The cell types clusters identified using the augmented signature matrix that was seeded with the 100 genes with the highest variance in the normal pancreas sample.

Combining the clustered cell types for the Top100 estimates increased the *ρ* to 0.32 but also increased the *RMSE* to 17.15. Similarly, the Augmented cell estimates had *ρ* = 0.58 and *RMSE* = 16.58. In other words, combining the cell types made it easier to get the relative order of cell type percentages correct, however the predicted fraction of cell types became less accurate.

### Hierarchical Clustering Improves Deconvolution Accuracy

**ADAPTS** Algorithm 3 (outlined in Hierarchical Deconvolution) was used to build custom signature matrices for breaking apart the clusters shown in Fig 5 and Fig 6. This improved deconvolution accuracies shown in the ‘hierarchical’ columns of Table 2. Applying the model built on the normal samples to the diabetic pancreas resulted in the even better blind predictive accuracies shown in Table 3 with the overall best accuracy provided by the hierarchical deconvolution using the Augmented signature matrix: *ρ* = 0.46, *RMSE*=8.91.

## Conclusion

Table 1 shows an example where using **ADAPTS** to include additional genes and tissue specific cell types improves the ability of a deconvolution algorithm to identify tumor fractions in microarray-based purified and mixed multiple myeloma gene expression samples. Thus we demonstrate that the techniques implemented in **ADAPTS** are potentially beneficial for many situations. The functions implemented in **ADAPTS** enable researchers to build their own custom signature matrices and investigate biosamples consisting of multiple cell types. Tables 2 and 3 show that these methods can build new signature matrices from single cell RNAseq (scRNAseq) data and effectively deconvolve the cell types determined by single cell analysis. This is expected to be particularly useful as researchers use scRNAseq to determine cell types that are present in tissue where large numbers of bulk gene expression samples are already available.

## Supporting information

**S1 Vig. ADAPTS.vignette.html**. ADAPTS (Automated Deconvolution Augmentation of Profiles for Tissue Specific cells) Vignette.

**S2 Vig. ADAPTS2.vignette.html**. ADAPTS Vignette 2: Single Cell Analysis.

## Acknowledgments

Thanks to Gareth Morgan, Jake Gockley, Robert Hershberg, Mary H Young, Andrew Dervan, and all other contributors to the paper “Identifying a High-risk Cellular Signature in the Multiple Myeloma Bone Marrow Microenvironment”.

