## Supplementary material for "ADAPTS: Automated Deconvolution Augmentation of Profiles for Tissue Specific cells": Vignette (from another version of this preprint: version 2, 2019-05-15)

#### Samuel Danziger, PhD

#### 2019-05-09

Part I - A: Deconvolve purified CD138+ tumor samples using LM22

```
library(ADAPTS)
library(ADAPTSdata)
library(ADAPTSdata2)
library(preprocessCore)
library(pheatmap)
doMC::registerDoMC(cores = parallel::detectCores())

LM22 <- ADAPTS::LM22
cd138p <- ADAPTSdata::cd138p
wbm <- ADAPTSdata2::wbm
tumorPer <- ADAPTSdata2::tumorPer
LM22.source <- ADAPTSdata2::fullLM22
addMGSM27 <- ADAPTSdata::addMGSM27
```

```
cellCounts <- estCellPercent.DCQ(LM22, cd138p)
tumorPer$LM22.CD138p.tumor <- colSums(cellCounts[c('Plasma.cells','B.cells.memory'),match(make.names(tumorPer$celfileRnas),colnames(cellCounts))])

incBool <- !is.na(tumorPer$CD138_Post_Sort) 
curData <- tumorPer[incBool,]
rmse.lm22.cd138p <- sqrt(mean((curData$CD138_Post_Sort - curData$LM22.CD138p.tumor)^2))
ct.lm22.cd138p <- cor.test(curData$CD138_Post_Sort, curData$LM22.CD138p.tumor)
titleStr <- paste0('LM22 Deconvolution of CD138+ Samples:\nRMSE = ', round(rmse.lm22.cd138p), '; Rho = ', round(ct.lm22.cd138p$estimate,2) )
plot(curData$CD138_Post_Sort, curData$LM22.CD138p.tumor, main=titleStr, xlab='% Purity', ylab='Deconvolved Tumor %')
```

Part I - B: Deconvolve WBM samples using LM22

```
cellCounts <- estCellPercent.DCQ(LM22, wbm)
tumorPer$LM22.WBM.tumor <- colSums(cellCounts[c('Plasma.cells','B.cells.memory'),match(make.names(tumorPer$celfileRnbx),colnames(cellCounts))])

tumorPer$PC_Mean <- rowMeans(tumorPer[,c('PC_AspNum','PC_BmbxNum')], na.rm=TRUE)
incBool <- !is.nan(tumorPer$PC_Mean) & !is.na(tumorPer$LM22.WBM.tumor)
curData <- tumorPer[incBool,] 
rmse.lm22.WBM <- sqrt(mean((curData$PC_Mean - curData$LM22.WBM.tumor)^2))
ct.lm22.WBM <- cor.test(curData$PC_Mean, curData$LM22.WBM.tumor)
titleStr <- paste0('LM22 Deconvolution of WBM Samples:\nRMSE = ', round(rmse.lm22.WBM), '; Rho = ', round(ct.lm22.WBM$estimate,2) )
plot(curData$PC_Mean, curData$LM22.WBM.tumor, main=titleStr, xlab='% WBM Tumor', ylab='Deconvolved Tumor %')
```

Part II - Build LM22 + ‘MM.plasma.cell’,‘PlasmaMemory’

```
LM22p <- LM22
augData <- addMGSM27[match(rownames(LM22), rownames(addMGSM27)), ]#grepl('PlasmaMemory',colnames(addMGSM27)) | grepl('MM.plasma.cell',colnames(addMGSM27))]
LM22p$PlasmaMemory <- rowSums(augData[,grepl('PlasmaMemory',colnames(augData))], na.rm=TRUE)
LM22p$MM.plasma.cell <- rowSums(augData[,grepl('MM.plasma.cell',colnames(augData))], na.rm=TRUE)
LM22p$osteoclast <- rowSums(augData[,grepl('osteoclast',colnames(augData))], na.rm=TRUE)
LM22p$osteoblast <- rowSums(augData[,grepl('osteoblast',colnames(augData))], na.rm=TRUE)
LM22p$adipocyte <- rowSums(augData[,grepl('adipocyte',colnames(augData))], na.rm=TRUE)
```

Part II - A: Deconvolve purified CD138+ tumor samples using LM22 + ‘MM.plasma.cell’,‘PlasmaMemory’

```
cellCounts <- estCellPercent.DCQ(LM22p, cd138p)
tumorPer$LM22p.CD138p.tumor <- colSums(cellCounts[c('Plasma.cells','B.cells.memory','MM.plasma.cell','PlasmaMemory'),match(make.names(tumorPer$celfileRnas),colnames(cellCounts))])

incBool <- !is.na(tumorPer$CD138_Post_Sort)
curData <- tumorPer[incBool,]
rmse.lm22p.cd138p <- sqrt(mean((curData$CD138_Post_Sort - curData$LM22p.CD138p.tumor)^2))
ct.lm22p.cd138p <- cor.test(curData$CD138_Post_Sort, curData$LM22p.CD138p.tumor)
titleStr <- paste0('LM22 + 5 Deconvolution of CD138+ Samples:\nRMSE = ', round(rmse.lm22p.cd138p), '; Rho = ', round(ct.lm22p.cd138p$estimate,2))
plot(curData$CD138_Post_Sort, curData$LM22p.CD138p.tumor, main=titleStr, xlab='% Purity', ylab='Deconvolved Tumor %')
```

Part II - B: Deconvolve WBM samples using LM22 + Bone Marrow Cells

```
cellCounts <- estCellPercent.DCQ(LM22p, wbm)
tumorPer$LM22p.WBM.tumor <- colSums(cellCounts[c('Plasma.cells','B.cells.memory','MM.plasma.cell','PlasmaMemory'),match(make.names(tumorPer$celfileRnbx),colnames(cellCounts))])

tumorPer$PC_Mean <- rowMeans(tumorPer[,c('PC_AspNum','PC_BmbxNum')], na.rm=TRUE)
incBool <- !is.nan(tumorPer$PC_Mean) & !is.na(tumorPer$LM22p.WBM.tumor)
curData <- tumorPer[incBool,] 
rmse.lm22p.WBM <- sqrt(mean((curData$PC_Mean - curData$LM22p.WBM.tumor)^2))
ct.lm22p.WBM <- cor.test(curData$PC_Mean, curData$LM22p.WBM.tumor)
titleStr <- paste0('LM22 + 5 Deconvolution of WBM Samples:\nRMSE = ', round(rmse.lm22p.WBM), '; Rho = ', round(ct.lm22p.WBM$estimate,2) )
plot(curData$PC_Mean, curData$LM22p.WBM.tumor, main=titleStr, xlab='% WBM Tumor', ylab='Deconvolved Tumor %')
```

Part III: Augmenting an existing signature matrix

```
olGenes <- rownames(LM22.source)[rownames(LM22.source) %in% rownames(addMGSM27)]
allData <- cbind(LM22.source[olGenes,], addMGSM27[olGenes,])
allData.nq <- as.data.frame(normalize.quantiles(as.matrix(allData)))
rownames(allData.nq) <- rownames(allData)
colnames(allData.nq) <- colnames(allData)
allData <- allData.nq

#Remove low variance genes to make the example run faster
allData.noNA <- allData[!apply(allData, 1, function(x){any(is.na(x))}),]
vars.noNA <- apply(allData.noNA, 1, var)

hist(vars.noNA)
```

```
#This will take a while.  To make it faster, filter genes by variance
subData <- allData[vars.noNA>2,]
gList <- rankByT(subData)
```

```
olgenes <- rownames(subData)

fullData <- allData[olgenes,colnames(LM22.source)]
colnames(fullData) <- sub('\\.[0-9]+$', '', colnames(fullData)) #Strip any trailing numbers added by make.names()

newData <- allData[olgenes,colnames(addMGSM27)]
colnames(newData) <- sub('\\.[0-9]+$', '', colnames(newData)) #Strip any trailing numbers added by make.names()

#MGSM27 <- AugmentSigMatrix(origMatrix=LM22, fullData=fullData, newData=newData, gList=gList, nGenes=1:100, plotToPDF=FALSE, imputeMissing=TRUE, condTol=1.01, postNorm=TRUE, minSumToRem=NA, addTitle=NULL, autoDetectMin=TRUE, calcSpillOver=FALSE)
#Note:  For the Identifying a High-risk Cellular Signature in the Multiple Myeloma Bone Marrow Microenvironment, we used the call commented out above, which resulted in 601 genes.  It looks like the package used to find the minimima has changed, results in a smaller number of genes (~570).  We recommend trying this option and looking at the condition number curves to see if it makes sense.  Future version will probably upgrade the 'autoDetectMin' option to be smarter. 

MGSM27 <- AugmentSigMatrix(origMatrix=LM22, fullData=fullData, newData=newData, gList=gList, nGenes=1:100, plotToPDF=FALSE, imputeMissing=TRUE, condTol=1.01, postNorm=TRUE, minSumToRem=NA, addTitle=NULL, autoDetectMin=FALSE, calcSpillOver=FALSE)
#>   missForest iteration 1 in progress...
#> randomForest 4.6-14
#> Type rfNews() to see new features/changes/bug fixes.
#> done!
#>   missForest iteration 2 in progress...done!
#>   missForest iteration 3 in progress...done!
#>   missForest iteration 4 in progress...done!
#>   missForest iteration 1 in progress...done!
#>   missForest iteration 2 in progress...done!
#>   missForest iteration 3 in progress...done!
#>   missForest iteration 4 in progress...done!
```

Part IV - A: Deconvolve purified CD138+ tumor samples using MGSM27

```
cellCounts <- estCellPercent.DCQ(MGSM27, cd138p)
tumorPer$MGSM27.CD138p.tumor <- colSums(cellCounts[c('Plasma.cells','B.cells.memory','MM.plasma.cell','PlasmaMemory'),match(make.names(tumorPer$celfileRnas),colnames(cellCounts))])

incBool <- !is.na(tumorPer$CD138_Post_Sort)
curData <- tumorPer[incBool,]
rmse.mgsm27.cd138p <- sqrt(mean((curData$CD138_Post_Sort - curData$MGSM27.CD138p.tumor)^2))
ct.mgsm27.cd138p <- cor.test(curData$CD138_Post_Sort, curData$MGSM27.CD138p.tumor)
titleStr <- paste0('MGSM27 Deconvolution of CD138+ Samples:\nRMSE = ', round(rmse.mgsm27.cd138p), '; Rho = ', round(ct.mgsm27.cd138p$estimate,2) )
plot(curData$CD138_Post_Sort, curData$MGSM27.CD138p.tumor, main=titleStr, xlab='% Purity', ylab='Deconvolved Tumor %')
```

Part IV - B: Deconvolve WBM samples using MGSM27

```
cellCounts <- estCellPercent.DCQ(MGSM27, wbm)
tumorPer$MGSM27.WBM.tumor <- colSums(cellCounts[c('Plasma.cells','B.cells.memory','MM.plasma.cell','PlasmaMemory'),match(make.names(tumorPer$celfileRnbx),colnames(cellCounts))])

tumorPer$PC_Mean <- rowMeans(tumorPer[,c('PC_AspNum','PC_BmbxNum')], na.rm=TRUE)
incBool <- !is.nan(tumorPer$PC_Mean) & !is.na(tumorPer$LM22.WBM.tumor)
curData <- tumorPer[incBool,] 
rmse.mgsm27.WBM <- sqrt(mean((as.numeric(curData$PC_Mean) - curData$MGSM27.WBM.tumor)^2))
ct.mgsm27.WBM <- cor.test(as.numeric(curData$PC_Mean), curData$MGSM27.WBM.tumor)
titleStr <- paste0('MGSM27 Deconvolution of WBM Samples:\nRMSE = ', round(rmse.mgsm27.WBM), '; Rho = ', round(ct.mgsm27.WBM$estimate,2) )
plot(curData$PC_Mean, curData$MGSM27.WBM.tumor, main=titleStr, xlab='% WBM Tumor', ylab='Deconvolved Tumor %')
```

Part V: Generating a new signature matrix (Top Variance, Subtract, Recreate)

```
olGenes <- rownames(LM22.source)[rownames(LM22.source) %in% rownames(addMGSM27)]
allData <- cbind(LM22.source[olGenes,], addMGSM27[olGenes,])
allData.nq <- as.data.frame(normalize.quantiles(as.matrix(allData)))
rownames(allData.nq) <- rownames(allData)
colnames(allData.nq) <- colnames(allData)
allData <- allData.nq
```

The top genes by variance will be the seedMatrix

```
nGenes <- 100

seedMat <- allData.noNA[names(tail(sort(vars.noNA), nGenes)),]
colnames(seedMat) <- sub('\\.[0-9]+$', '', colnames(seedMat))
seedMat.sig <- lapply(unique(colnames(seedMat)), function(x){ rowMeans(seedMat[,colnames(seedMat)==x], na.rm=TRUE)})
seedMat.sig <- do.call(cbind, seedMat.sig)
colnames(seedMat.sig) <- unique(colnames(seedMat))
pheatmap(seedMat.sig)
```

Augment that top gene network

```
subData.noNA <- allData.noNA[vars.noNA>2,]
colnames(allData.noNA) <- sub('\\.[0-9]+$', '', colnames(allData.noNA))
gList.noNA <- rankByT(subData.noNA)
```

```
topAug <- AugmentSigMatrix(origMatrix=seedMat.sig, fullData=allData.noNA, newData=allData.noNA, gList=gList.noNA, nGenes=1:100, plotToPDF=FALSE, imputeMissing=TRUE, condTol=1.01, postNorm=TRUE, minSumToRem=NA, addTitle=NULL, autoDetectMin=FALSE, calcSpillOver=FALSE)
#>   missForest iteration 1 in progress...done!
#>   missForest iteration 2 in progress...done!
#>   missForest iteration 1 in progress...done!
#>   missForest iteration 2 in progress...done!
```

Remove the seed genes and then rebuild

```
topAug.remSeed <- topAug[!(rownames(topAug) %in% rownames(seedMat.sig)),]

topAug2 <- AugmentSigMatrix(origMatrix=topAug.remSeed, fullData=allData.noNA, newData=allData.noNA, gList=gList.noNA, nGenes=1:100, plotToPDF=FALSE, imputeMissing=TRUE, condTol=1.01, postNorm=TRUE, minSumToRem=NA, addTitle=NULL, autoDetectMin=FALSE, calcSpillOver=FALSE)
#>   missForest iteration 1 in progress...done!
#>   missForest iteration 2 in progress...done!
#>   missForest iteration 1 in progress...done!
#>   missForest iteration 2 in progress...done!
```

Part VI - A: Deconvolve purified CD138+ tumor samples using de-novo deconvolution matrix

```
cellCounts <- estCellPercent.DCQ(topAug2, cd138p)
tumorPer$DeNovo.CD138p.tumor <- colSums(cellCounts[c('Plasma.cells','B.cells.memory','MM.plasma.cell','PlasmaMemory'),match(make.names(tumorPer$celfileRnas),colnames(cellCounts))])

incBool <- !is.na(tumorPer$CD138_Post_Sort)
curData <- tumorPer[incBool,]
rmse.DeNovo.cd138p <- sqrt(mean((curData$CD138_Post_Sort - curData$DeNovo.CD138p.tumor)^2))
ct.DeNovo.cd138p <- cor.test(curData$CD138_Post_Sort, curData$DeNovo.CD138p.tumor)
titleStr <- paste0('DeNovo Deconvolution of CD138+ Samples:\nRMSE = ', round(rmse.DeNovo.cd138p), '; Rho = ', round(ct.DeNovo.cd138p$estimate,2) )
plot(curData$CD138_Post_Sort, curData$DeNovo.CD138p.tumor, main=titleStr, xlab='% Purity', ylab='Deconvolved Tumor %')
```

Part VI - B: Deconvolve WBM samples using de-novo deconvolution matrix

```
cellCounts <- estCellPercent.DCQ(topAug2, wbm)
tumorPer$DeNovo.WBM.tumor <- colSums(cellCounts[c('Plasma.cells','B.cells.memory','MM.plasma.cell','PlasmaMemory'),match(make.names(tumorPer$celfileRnbx),colnames(cellCounts))])

tumorPer$PC_Mean <- rowMeans(tumorPer[,c('PC_AspNum','PC_BmbxNum')], na.rm=TRUE)
incBool <- !is.nan(tumorPer$PC_Mean) & !is.na(tumorPer$DeNovo.WBM.tumor)
curData <- tumorPer[incBool,] 
rmse.DeNovo.WBM <- sqrt(mean((curData$PC_Mean - curData$DeNovo.WBM.tumor)^2))
ct.DeNovo.WBM <- cor.test(curData$PC_Mean, curData$DeNovo.WBM.tumor)
titleStr <- paste0('DeNovo Deconvolution of WBM Samples:\nRMSE = ', round(rmse.DeNovo.WBM), '; Rho = ', round(ct.DeNovo.WBM$estimate,2) )
plot(curData$PC_Mean, curData$DeNovo.WBM.tumor, main=titleStr, xlab='% WBM Tumor', ylab='Deconvolved Tumor %')
```

Build the summary table

```
outMatrix <- matrix(0, nrow=4, ncol=4, dimnames=list(c('LM22','LM22 + 5','MGSM27','DeNovo MGSM27'),c('CD138+.RMSE','CD138+.Rho','WBM.RMSE','WBM.Rho')))
outMatrix[1,1] <- rmse.lm22.cd138p
outMatrix[1,2] <- ct.lm22.cd138p$estimate
outMatrix[1,3] <- rmse.lm22.WBM
outMatrix[1,4] <- ct.lm22.WBM$estimate

outMatrix[2,1] <- rmse.lm22p.cd138p
outMatrix[2,2] <- ct.lm22p.cd138p$estimate
outMatrix[2,3] <- rmse.lm22p.WBM
outMatrix[2,4] <- ct.lm22p.WBM$estimate
  
outMatrix[3,1] <- rmse.mgsm27.cd138p
outMatrix[3,2] <- ct.mgsm27.cd138p$estimate
outMatrix[3,3] <- rmse.mgsm27.WBM
outMatrix[3,4] <- ct.mgsm27.WBM$estimate

outMatrix[4,1] <- rmse.DeNovo.cd138p
outMatrix[4,2] <- ct.DeNovo.cd138p$estimate
outMatrix[4,3] <- rmse.DeNovo.WBM
outMatrix[4,4] <- ct.DeNovo.WBM$estimate

print(signif(outMatrix,2))
#>               CD138+.RMSE CD138+.Rho WBM.RMSE WBM.Rho
#> LM22                 27.0       0.26       38    0.59
#> LM22 + 5             12.0       0.26       37    0.53
#> MGSM27                8.8       0.33       23    0.60
#> DeNovo MGSM27        36.0       0.18       24    0.65
```
