## Supplementary material for "ADAPTS: Automated Deconvolution Augmentation of Profiles for Tissue Specific cells": R package: ADAPTS.vignette.html (from another version of this preprint: version 2, 2019-05-15)

#### Samuel Danziger, PhD

#### 2019-05-14

Part I - A: Deconvolve purified CD138+ tumor samples using LM22 Note: ADAPTSdata and ADAPTSdata2 come from GitHub devtools::install\_github(“sdanzige/ADAPTSdata”) devtools::install\_github(“sdanzige/ADAPTSdata2”)

```
#This will take a while.  To make it faster, filter genes by variance
subData <- allData[vars.noNA>2,]
gList <- rankByT(subData)
```

```
olgenes <- rownames(subData)

fullData <- allData[olgenes,colnames(LM22.source)]
colnames(fullData) <- sub('\\.[0-9]+$', '', colnames(fullData)) #Strip any trailing numbers added by make.names()

newData <- allData[olgenes,colnames(addMGSM27)]
colnames(newData) <- sub('\\.[0-9]+$', '', colnames(newData)) #Strip any trailing numbers added by make.names()

#MGSM27 <- AugmentSigMatrix(origMatrix=LM22, fullData=fullData, newData=newData, gList=gList, nGenes=1:100, plotToPDF=FALSE, imputeMissing=TRUE, condTol=1.01, postNorm=TRUE, minSumToRem=NA, addTitle=NULL, autoDetectMin=TRUE, calcSpillOver=FALSE)
#Note:  For the Identifying a High-risk Cellular Signature in the Multiple Myeloma Bone Marrow Microenvironment, we used the call commented out above, which resulted in 601 genes.  It looks like the package used to find the minimima has changed, results in a smaller number of genes (~570).  We recommend trying this option and looking at the condition number curves to see if it makes sense.  Future version will probably upgrade the 'autoDetectMin' option to be smarter. 

MGSM27 <- AugmentSigMatrix(origMatrix=LM22, fullData=fullData, newData=newData, gList=gList, nGenes=1:100, plotToPDF=FALSE, imputeMissing=TRUE, condTol=1.01, postNorm=TRUE, minSumToRem=NA, addTitle=NULL, autoDetectMin=FALSE, calcSpillOver=FALSE)
#>   missForest iteration 1 in progress...
#> randomForest 4.6-14
#> Type rfNews() to see new features/changes/bug fixes.
#> done!
#>   missForest iteration 2 in progress...done!
#>   missForest iteration 3 in progress...done!
#>   missForest iteration 4 in progress...done!
#>   missForest iteration 5 in progress...done!
#>   missForest iteration 6 in progress...done!
#>   missForest iteration 7 in progress...done!
#>   missForest iteration 1 in progress...done!
#>   missForest iteration 2 in progress...done!
#>   missForest iteration 3 in progress...done!
#>   missForest iteration 4 in progress...done!
```

Build the summary table

```
outMatrix <- matrix(0, nrow=4, ncol=4, dimnames=list(c('LM22','LM22 + 5','MGSM27','DeNovo MGSM27'),c('CD138+.RMSE','CD138+.Rho','WBM.RMSE','WBM.Rho')))
outMatrix[1,1] <- rmse.lm22.cd138p
outMatrix[1,2] <- ct.lm22.cd138p$estimate
outMatrix[1,3] <- rmse.lm22.WBM
outMatrix[1,4] <- ct.lm22.WBM$estimate

outMatrix[2,1] <- rmse.lm22p.cd138p
outMatrix[2,2] <- ct.lm22p.cd138p$estimate
outMatrix[2,3] <- rmse.lm22p.WBM
outMatrix[2,4] <- ct.lm22p.WBM$estimate
  
outMatrix[3,1] <- rmse.mgsm27.cd138p
outMatrix[3,2] <- ct.mgsm27.cd138p$estimate
outMatrix[3,3] <- rmse.mgsm27.WBM
outMatrix[3,4] <- ct.mgsm27.WBM$estimate

outMatrix[4,1] <- rmse.DeNovo.cd138p
outMatrix[4,2] <- ct.DeNovo.cd138p$estimate
outMatrix[4,3] <- rmse.DeNovo.WBM
outMatrix[4,4] <- ct.DeNovo.WBM$estimate

print(signif(outMatrix,2))
#>               CD138+.RMSE CD138+.Rho WBM.RMSE WBM.Rho
#> LM22                 27.0       0.26       38    0.59
#> LM22 + 5             12.0       0.26       37    0.53
#> MGSM27                8.7       0.33       23    0.60
#> DeNovo MGSM27        36.0       0.18       24    0.65
```
